# DNA damage in 3D constricted migration or after lamin-A depletion in 2D: shared mechanisms of repair factor mis-localization under nuclear stress

**DOI:** 10.1101/097162

**Authors:** Yuntao Xia, Jerome Irianto, Charlotte R. Pfeifer, Jiazheng Ji, Irena L. Ivanovska, Manu Tewari, Rachel R. Bennett, Shane M. Harding, Andrea J. Liu, Roger A. Greenberg, Dennis E. Discher

**Affiliations:** Physical Sciences Oncology Center at Penn (PSOC@Penn), 129 Towne Bldg., University of Pennsylvania, Philadelphia, PA 19104, USA; Molecular & Cell Biophysics Lab, 129 Towne Bldg., University of Pennsylvania, Philadelphia, PA 19104, USA; Graduate Group / Department of Physics & Astronomy, 129 Towne Bldg., University of Pennsylvania, Philadelphia, PA 19104, USA; Cancer Biology, Abramson Family Cancer Research Institute, Perelman School of Medicine, 129 Towne Bldg., University of Pennsylvania, Philadelphia, PA 19104, USA

## Abstract

Cells that migrate through small, rigid pores and that have normal levels of the nuclear structure protein lamin-A exhibit an increase in DNA damage, which is also observed with lamin-A depletion in diseases such as cancer and with many lamin-A mutations. Here we show nuclear envelope rupture is a shared feature that increases in standard culture after lamin-A knockdown, which causes nuclear loss of multiple DNA repair factors and increased DNA damage. Some repair factors are merely mis-localized to cytoplasm whereas others are partially depleted unless rescued by lamin-A expression. Compared to standard cultures on rigid glass coverslips, the growth of lamin-A low cells on soft matrix relaxes cytoskeletal stress on the nucleus, suppresses the mis-localization of DNA repair factors, and minimizes DNA damage nearly to wildtype levels. Conversely, constricted migration of the lamin-A low cells causes abnormally high levels of DNA damage, consistent with sustained loss of repair factors. The findings add insight into why monogenic progeroid syndromes that often associate with increased DNA damage and predominantly impact cells in stiff tissues result from mutations only in lamin-A or DNA repair factors.

## Introduction

A recent report of migration-induced DNA damage based on loss of multiple repair factor(s)^10^ seems similar in mechanism to disparate studies of lamin-A null and mutant fibroblasts in standard culture^7,5^. For cultures of at least one mutant, repetitive rupture of the nuclear envelope (NE) in cells cultured on rigid surfaces can be minimized by soft, tissue-like substrates that reduce nuclear stress^19^, and the rupture causes cytoplasmic mis-localization of at least a few mobile nuclear factors^5^. Mis-localization after rupture should in principle include many mobile DNA repair factors^9^, but these have not been imaged after lamin-A mutation or depletion^5,7,19^. In particular, the mobile repair factor 53BP1 might be expected to be more cytoplasmic after rupture, which could explain its rapid degradation in lamin-A null fibroblasts (in the absence of any transcript change) as well as the observation that overexpression of 53BP1 can rescue irradiation-induced DNA damage in lamin-A low cells^7^. Loss of 53BP1 does occur early across many human cancers of different tissue and cell types, but its loss decouples from the appearance of the DNA damage marker γH2AX^15^. Furthermore, overexpression of 53BP1 does not reproducibly provide a significant rescue of DNA damage after constricted migration of U2OS osteosarcoma cells that express abundant lamin-A^10^.

For other repair factors that regulate chromosome copy number variations – including BRCA's that explain genomic aberrations in osteosarcoma^12^, mis-localization after constricted migration^10^ could explain some of the genomic variation that results from migration of U2OS cells through rigid 3 μm pores more so than 8 μm pores^10^. The fact that lamin-A is highly mutated but does not increase the risk for cancer (unlike mutations in repair factors such as BRCA’s) further implicates other important mechanisms unrelated to rupture, such as squeezing-dependent segregation of repair factors away from chromatin^11^. Furthermore, while an EMT-like change in cancer cell phenotype after constricted migration^10^ has illustrated invasion-mutation mechanisms pertinent to metastasis and to the heterogeneity within and between tumors^6^, mechanistic links between the seminal observations of DNA damage, repair factor depletion, and genome variation^10^ could benefit from more direct comparisons with the effects of Lamin-A depletion combined with a few key rescue experiments.

Additional motivation for studying Lamin-A and repair factor dynamics is provided by the well-characterized Progeroid Syndromes that are monogenic disorders resulting either from mutations in DNA repair factors or else from select mutations in Lamin-A. The accelerated aging that results affects a majority of tissues, especially stiff tissues such as bone, muscle, and skin; Progeroid Syndromes do not result from mutations in nucleases or oxidation/metabolism factors implicated in DNA damage in other contexts. The RecQ helicase WRN is functional, for example, when interacting with DNA in the nucleus, but mis-localization of WRN in the cytoplasm as associated with the Progeroid Syndrome known as Werner syndrome causes as increase in protein degradation in multiple cell types (including U2OS cells) within 16 hours^17^. A higher risk of cancer with WRN mutations is generally evident with Progeroid Syndromes involving DNA repair factor mutations, such as those in RECQL4 that increase the risk of osteosarcoma^13^. Genomic alterations, DNA damage, aging and cancer are thus strongly linked, and mis-localization of repair factors during NE rupture could contribute. Our principle hypothesis here is that nuclear stress can cause NE rupture leading to repair factor loss and DNA damage in low lamin-A cells in conventional 2D cultures on rigid substrates, with similar mechanisms for a large DNA damage increase in lamin-A low cells that have squeezed through small 3D pores.

## Results and Discussion

### Mis-localization of 53BP1 in constricted migration and also in cells on rigid glass

U2OS osteosarcoma cells from the stiff matrix of bones express high levels of lamin-A, relative to cells from softer tissue^8^, but U2OS cells can still squeeze through a rigid transwell filter with 3-μm pores even in the absence of any serum gradient^9,10^. During this constricted migration, the nucleus is under sufficiently high stress to rupture lamin-B and the NE with release of mobile nuclear factors including DNA repair factors Ku80 and BRCA1^9,10^(Fig 1A); and 53BP1 is one more repair factor that exhibits a decreased Nuc/Cyto intensity ratio after migration (Fig 1B,C). Increased DNA breaks after pore migration are evident as damage foci of γH2AX that are randomly distributed within the nucleoplasm (Fig 1B), consistent with the distribution of foci after partial knockdown of DNA repair factors Ku80 and BRCA1 in 2D cultures of U2OS cells^10^.

**Fig. 1.**
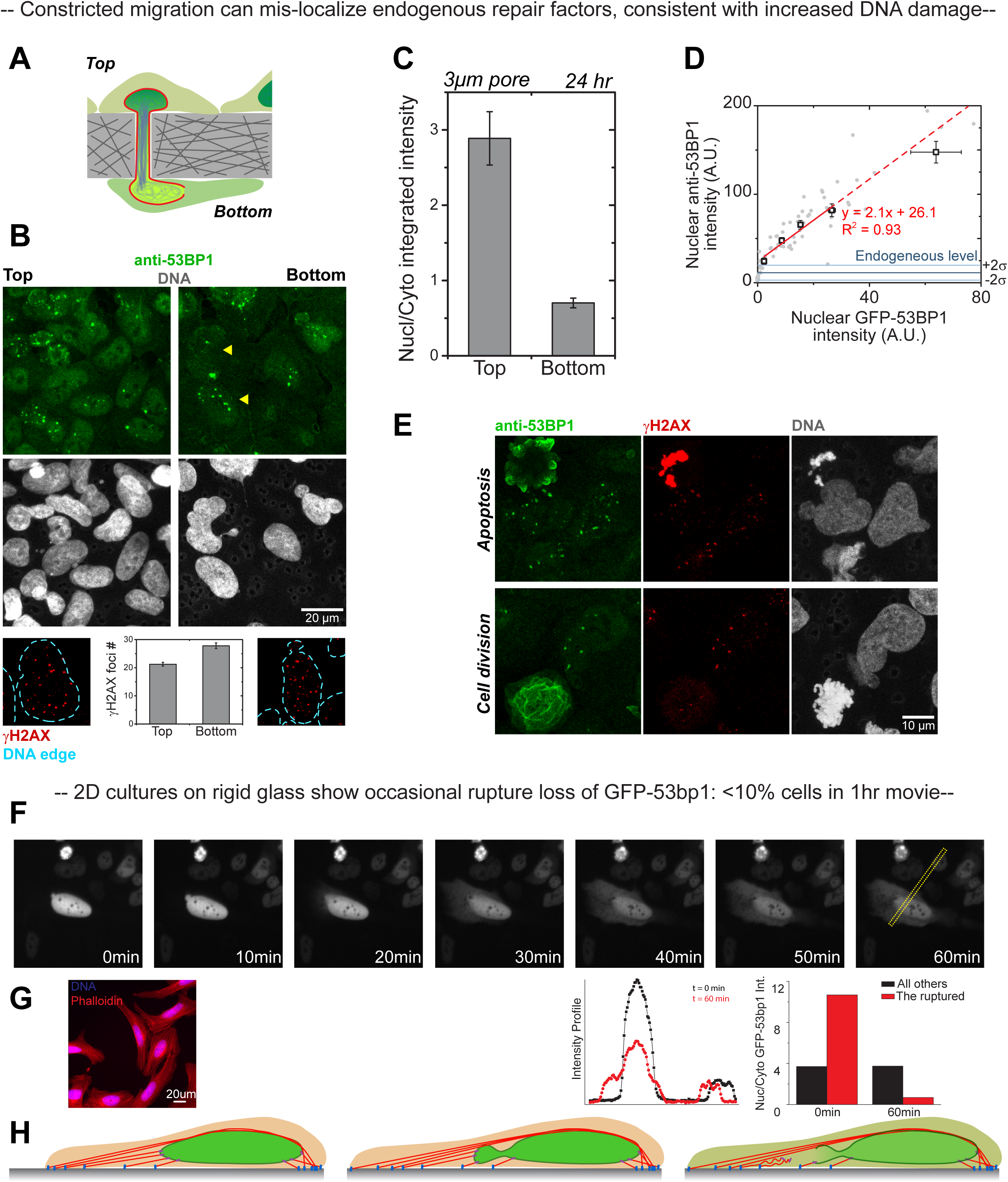
Nuclear envelop rupture and mis-localization of repair factors are observed in both 3D constricted migration and 2D culture on rigid glass. (A) Schematics illustrate that as a cell nucleus squeezes into a small pore, its chromatin becomes compacts and both pore exclusion and lamin-B rupture can occur. The green indicates nuclear diffusible protein and the red indicates nuclear lamin-B. (B) U2OS cells on both sides of the filter contain 53BP1 protein. Increased DNA DSBs are observed post-migration μm) based on immunostaining of γH2AX and these DSBs are evenly distributed in the nucleus. (>100 cells are quantified per condition, n>5 experiments, average ± SEM.) (C) For 53BP1, nuclear-to-cytoplasmic intensity ratio decreases in U2OS cells post-migration. (>10 fields of view per condition, average ± SEM.) (D) Specificity of 53BP1 antibody was validated by immuno-staining U2OS cells with GFP-53BP1 over-expression. At lower GFP intensity level, 53BP1 antibody intensity is statistically the same as the non-transfected cells. At higher over-expression level, 53BP1 antibody intensity increases proportionally to GFP intensity. (E) Cells during apoptosis shows cytoplasmic 53BP1, and bright γH2AX co-localized with DNA. Cells during mitosis show cytoplasmic 53BP1 as well but no bright γH2AX is seen. (F) U2OS cells on rigid glass show occasional rupture loss of GFP-53BP1 within 1hr. NE rupture causes nuclear depletion of 53BP1, consistent with drop of Nuc/Cyto 53BP1 intensity ratio. (G) U2OS cells have many long stress fibers. (H) Schematics illustrate nuclear bleb formation and rupture on rigid glass.

The change in 53BP1 Nuc/Cyto ratio is assessed at the single cell level with an antibody against endogenous 53BP1 that gives a linear increase in ‘red’ fluorescence in cells expressing increasing levels of green GFP-53BP1 (Fig 1D). 53BP1 is a repair factor with specific roles in DNA repair^2^, that could explain why it does not always co-localize with γH2AX (eg. apoptosis or mitosis: Fig 1E), but the pertinent observation here is that this nuclear factor mis-localizes after constricted migration.

The nucleus of a cell that is well-spread on 2D rigid glass is also under moderate stress as exerted by the cytoskeleton^1,18^. Nuclear rupture can be occasionally observed and exhibits at least some features in common with pore migration: an extruded bleb of chromatin forms followed by evident rupture with loss of a mobile repair factor such as GFP-53BP1 (Fig 1F). A very large increase in total cytoplasmic signal is accompanied by a modest decrease in total nuclear signal (Fig 1F, **plots**). Well-spread U2OS cells have the expected abundance of stress fibers (Fig 1G) that can apply high forces to a rigid substrate and to thereby sculpt nuclear shape (Fig 1H)^1,18^ The low fraction of wildtype cells that rupture during such a movie (<10%) is much lower than after constricted migration, consistent with much higher nuclear stress in migration and also consistent with observations of wildtype fibroblasts in studies of lamin-A mutations and deficiencies^7,5^. However, lamin-A defective cells show 5-10 fold more frequent nuclear rupture than wildtype cells with rupture occurring every 30 min in lamin-A nulls^5, 19^. Proteins that have been seen to mis-localize have thus far included transcription factors (OCT1, PML and Rel A) with effects on downstream gene expression^5^. Mis-localization of DNA repair factors has not yet been examined in lamin-A compromised cells.

### Endogenous repair factors are more cytoplasmic after partial lamin-A knockdown

To assess the role of Lamin-A in nuclear envelope rupture, stable knockdown with shLMNA and appropriate rescues were achieved with A549 lung carcinoma cells. A similarly stable knockdown could not be achieved in the U2OS line, which hints at the potential importance of nuclear integrity. Indeed, A549 knockdown cells – with ~30% of lamin-A relative to control cells – have been shown in micropipette aspiration to have decreased nuclear stiffness and to undergo nuclear rupture at much lower stress^9^. The nucleus of A549 cells is also more spread and flattened in cultures on stiff gels than on soft gels^1,18^, consistent with stiffness-driven nuclear stress. Importantly, the mobile nucleoplasmic repair factor Ku80 is prominent in the cytoplasm of ~10 fold more cells in knockdown cultures than controls (Fig 2A,B). Antibody specificity for immunofluorescence is validated by a decreased signal after partial knockdown with si-Ku80 (Fig 2B, **small bargraph**) and previously by a linear increase in signal with GFP-Ku80 expression^10^.

**Fig. 2.**
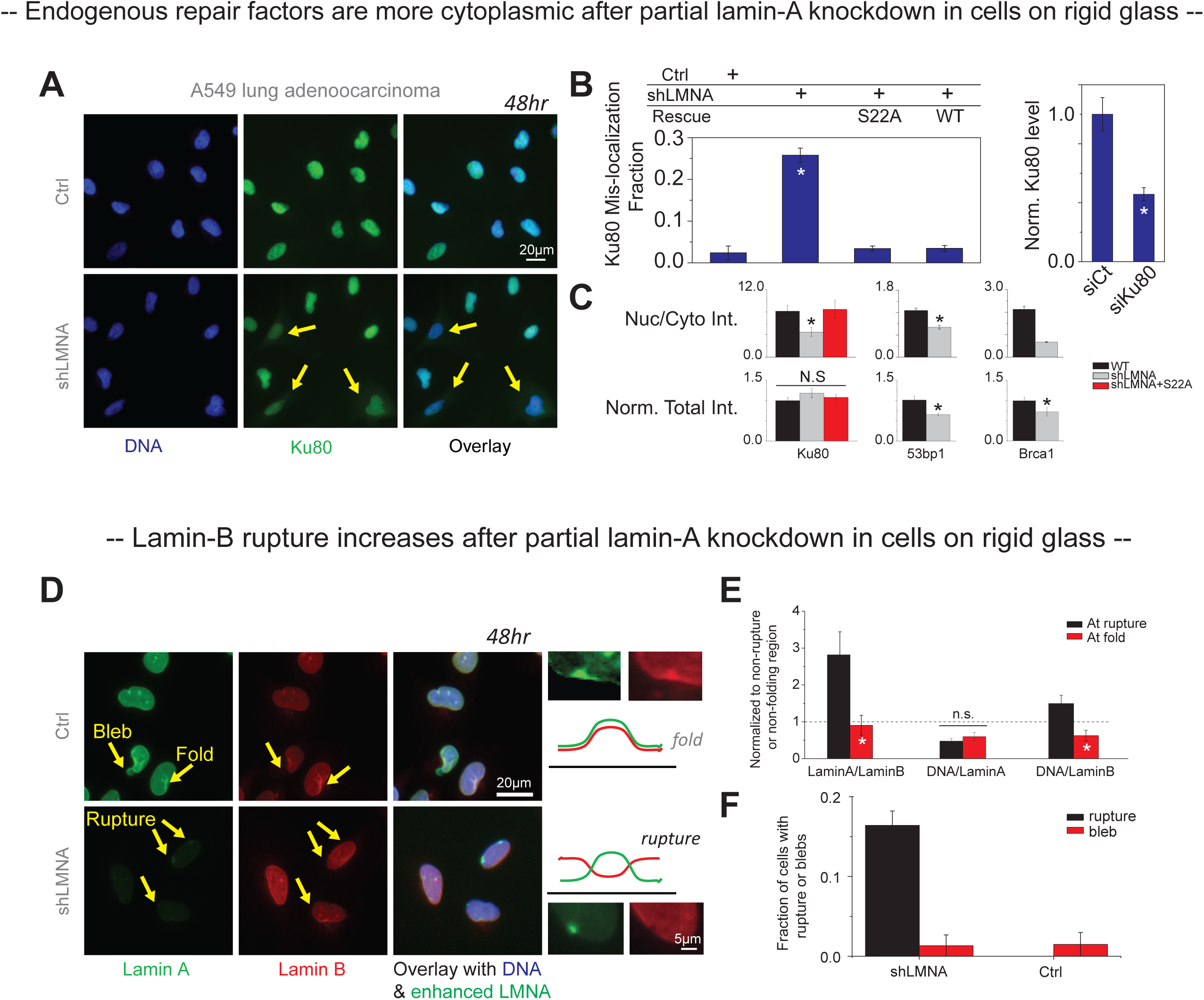
Mis-localization of endogenous repair factors and lamin-B rupture appear more frequent after partial lamin-A knockdown in cells on rigid glass. (A) Immunostaining of repair factor Ku80 highlight nuclear localization for Ctrl cells cultured on rigid glass (48hr), but cytoplasmic mis-localization after partial lamin-A knockdown. Arrows point to the cytoplasmic signal of Ku80. (B) 25.7% of shLMNA cells on rigid glass show Ku80 mis-localization comparing to 2.4% in Ctrl cells, and fraction of cells with Ku80 mis-localization can be efficiently rescued by expression of non-phosphrylatable, stabilizing mutant S22A lamin-A or wildtype lamin-A. Antibody specificity was validated by immunostaining of siCtrl and siKu80 samples. (>200 cells per condition, average ± SEM. *P<0.05 to Ctrl) (C) For Ku80, 53BP1 and BRCA1, nuclear-to-cytoplasmic intensity ratio decreases in shLMNA cells, indicating mis-localization. Ctrl and S22A rescued cells show no difference in Nuc/Cyto ratio for Ku80, indicating mis-localization is rescued. Total intensity of Ku80 per cell in Ctrl, shLMNA and S22A rescued cells does not change, but 53BP1 and BRCA1 total intensity per cell drops in shLMNA cells. (>5 fields of view per condition, average ± SEM. *P<0.05 to Ctrl) (D) Blebs in Ctrl cells and lamin-B ruptures in shLMNA cells are enriched in lamin-A but lack of lamin B, whereas fold sites are enriched in both lamin-A and lamin-B. (E) The ratios of Lamin-A/lamin-B, DNA/lamin-A, DNA/lamin-B are normalized to non-ruptured or non-folding region. *P<0.05 to Ctrl) (F) Fraction of cells with ruptures or blebs for shLMNA and Ctrl cells are counted based on (D). (>5 fields of view per condition, average ± SEM.)

For rescue of the knockdown phenotype, both wildtype Lamin-A and a lamina-stabilizing, non-phosphorylatable construct S22A were tested as they are known to exert similar effects on mechanosensing in the A549 cells^1,18^. Expression of both constructs dramatically decrease the fraction of cells with cytoplasmic Ku80 to control levels (Fig 2B) and thereby increase the Nuc/Cyto ratio without affecting total Ku80 intensities (Fig. 2C). Both 53BP1 and BRCA1 also show decreased Nuc/Cyto ratios in knockdown cells while also exhibiting decreased total intensities (Fig 2C). The constant overall level of Ku80 makes it a better marker of mis-localization, but the results for all three repair factors prove consistent with NE rupture and prompted a more direct visualization of the nuclear lamina.

As suggested by Fig 1H and by past studies of wildtype cells cultured on rigid substrates^7,5^ or in constricted migration^10^, lamina-disrupted blebs can form in wildtype nuclei, but only ~1-2% of A549 Ctrl cells exhibit such blebs (Fig 2D). In the cells with partial knockdown of lamin-A, lamina disruption is evident at the nuclear rim with a strong decrease in lamin B coincident with a strong increase in the residual Lamin-A (Fig 2D, E). The nuclear rim is where strain in the lamina is estimated to be greatest in strongly spread cells^1^. Wrinkles or folds are also evident in the NE, but folds are distinct in being enriched in both types of lamins. Based on the anti-correlated signature for NE rupture, about 10-fold more knockdown cells exhibit lamina disruptions or blebs versus control cells (Fig 2F). The fold difference is consistent with the increased fraction of cells exhibiting mis-localization of repair factor (Fig 2B).

### Repair factor levels selectively affected by lamin-A have different effects on DNA damage

Loss of a large fraction of BRCA1 and 53BP1 has been reported for lamin-A null fibroblasts in standard cultures that nonetheless maintain normal levels of Ku80^16^. Partial depletion of lamin-A has similar effects with A549 cells: immunoblots show 53BP1 is partially lost in shLMNA cells and is rescued in clones expressing wildtype and S22A lamin-A (Fig 3A), whereas Ku80 is unaffected (Fig 3B). BRCA1 is also partially lost from the shLMNA cells (Fig 3C), and since past studies showed *BRCA1* message is decreased in level upon Lamin-A depletion (unlike 53BP1) ^4^, *BRCA1* message was directly knocked down in wildtype cells to assess an effect on Ku80 protein levels – which do not change (Fig 3D). ChIP-Seq has shown BRCA1 protein binds the promoter regions of many genes^14^, as does the SRF transcription factor that decreases in Lamin-A knockdown cells^1^; both BRCA1 and SRF bind the promoter region of *BRCA1* (Fig 3E) but not that of Ku80 (*XRCC5* gene). Further studies of mechanism are of course required, but the results are consistent with past studies and immunofluorescence here (Fig 2C) in documenting the selective loss of DNA repair factors.

**Fig. 3.**
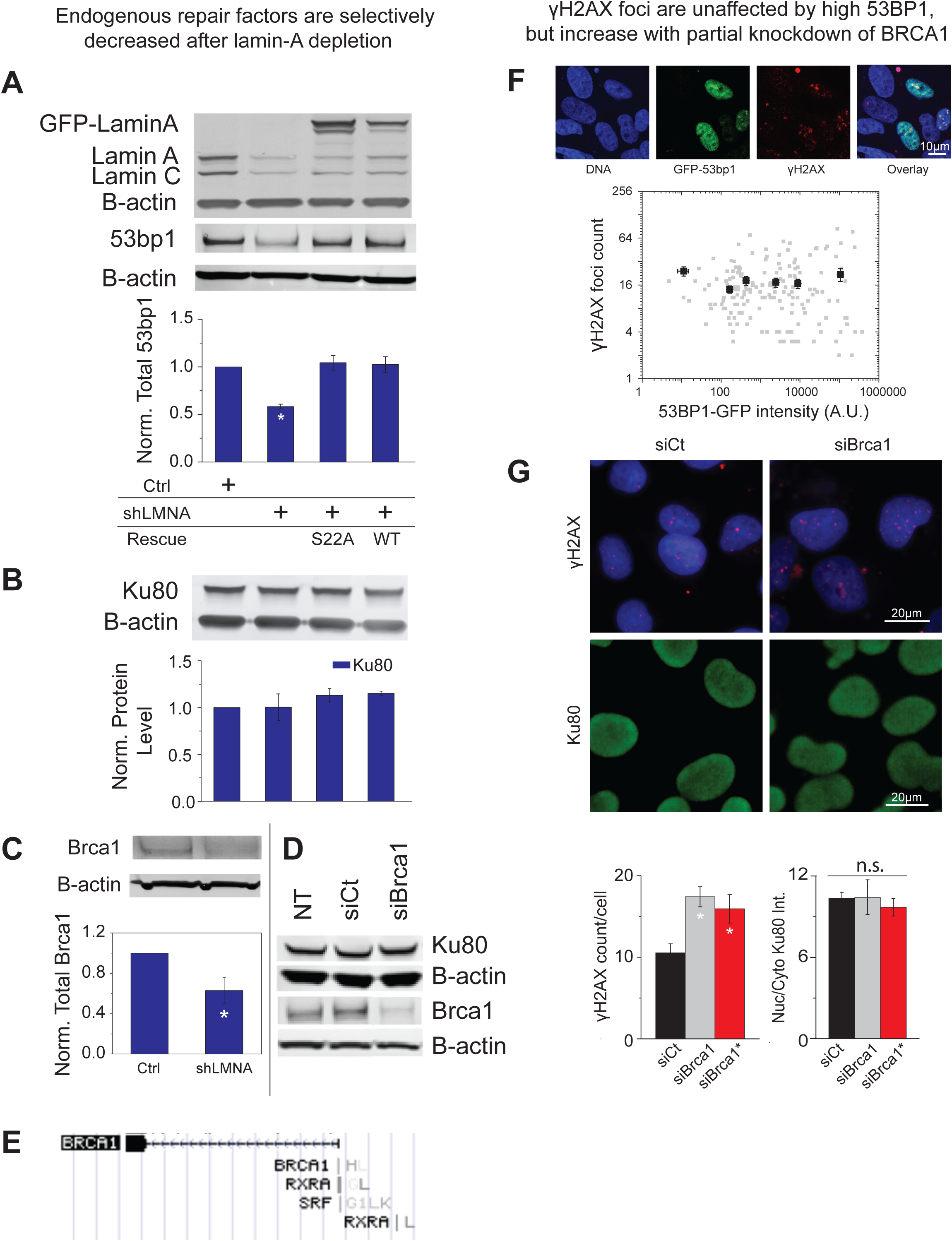
Endogenous repair factors are selectively decreased after lamin-A depletion and BRCA1 rather than 53BP1 shifts DNA damage/repair equilibrium. (A) Western blot is used to confirm LMNA knockdown or rescue. shLMNA cells have lower level of 53BP1 protein. (>3 western blots per condition, n ≥ 2 experiments, average ± SEM. *P<0.05 to Ctrl) (B) Western blot shows Ku80 protein level is constant for all Ctrl, shLMNA and Rescued cells. (>3 western blots per condition, n ≥ 2 experiments, average ± SEM.) (C) Western blot shows shLMNA cells have lower level of BRCA1 comparing to Ctrl cells. (>3 western blots per condition, n ≥ 2 experiments, average ± SEM. *P<0.05 to Ctrl) (D) siBRCA1 specifically knockdowns BRCA1, and Ku80 or B-actin is unaffected. (>3 western blots per condition, n ≥ 2 experiments, average ± SEM.) (E) BRCA1, RXRA and SRF can bind to the promoter region of BRCA1. (From Encode) (F) Overexpression of GFP-53BP1 does not reproducibly decrease γH2AX foci number. (G) 72 hr transient depletion of BRCA1 results in more γH2AX foci but no significant difference of Ku80 mis-localization is observed. (≥ 150 nuclei per conditions, n = 2 experiments, average ± SEM. *P<0.05 to Ctrl)

Functional effects of altered levels of DNA repair factors on DNA damage were assessed in terms of basal counts of γH2AX foci. Transfection of U2OS cells with GFP-53BP1 does not reproducibly indicate a significant effect on γH2AX foci counts (Fig 3F), with similar experiments on A549 cells causing cell death. Transfection of U2OS cells with si-BRCA1, however, did increase γH2AX foci counts and also showed no effect on nuclear envelope rupture based on Ku80 localization (Fig 3G). Importantly, the nucleoplasmic distribution of γH2AX foci after constricted migration is also evident after BRCA1 knockdown; in neither case is there an obvious localization of large or numerous foci near the nuclear edge where rupture might have occurred (Fig 1F, 2D). The increase in DNA damage with BRCA1 knockdown is therefore attributable to a decrease in the level of this mobile repair factor that diffuses to dispersed sites of damage (eg. sites of damaged synthesis^2^). The results further indicate that at least this one effector which is downstream of rupture does not propagate back upstream.

### DNA damage after lamin-A knockdown is rescued by soft matrix, which decreases rupture

Depletion of repair factors by either siRNAs or rupture has been linked to increased DNA damage^10^. Likewise, the shLMNA cells here exhibit more γH2AX foci than Ctrl cells under standard conditions of culture on rigid plastic (Fig 4A), and once again the γH2AX foci are distributed in the nucleoplasm rather than appearing concentrated at the nuclear envelope. Moreover, DNA damage in shLMNA cells can be rescued by expression of either the wildtype Lamin-A or the S22A stable form of Lamin-A (Fig 4A). The result is consistent with rescue of both the rupture-induced mis-localization of repair factors and their depletion (Fig 2B, 3A).

**Fig. 4.**
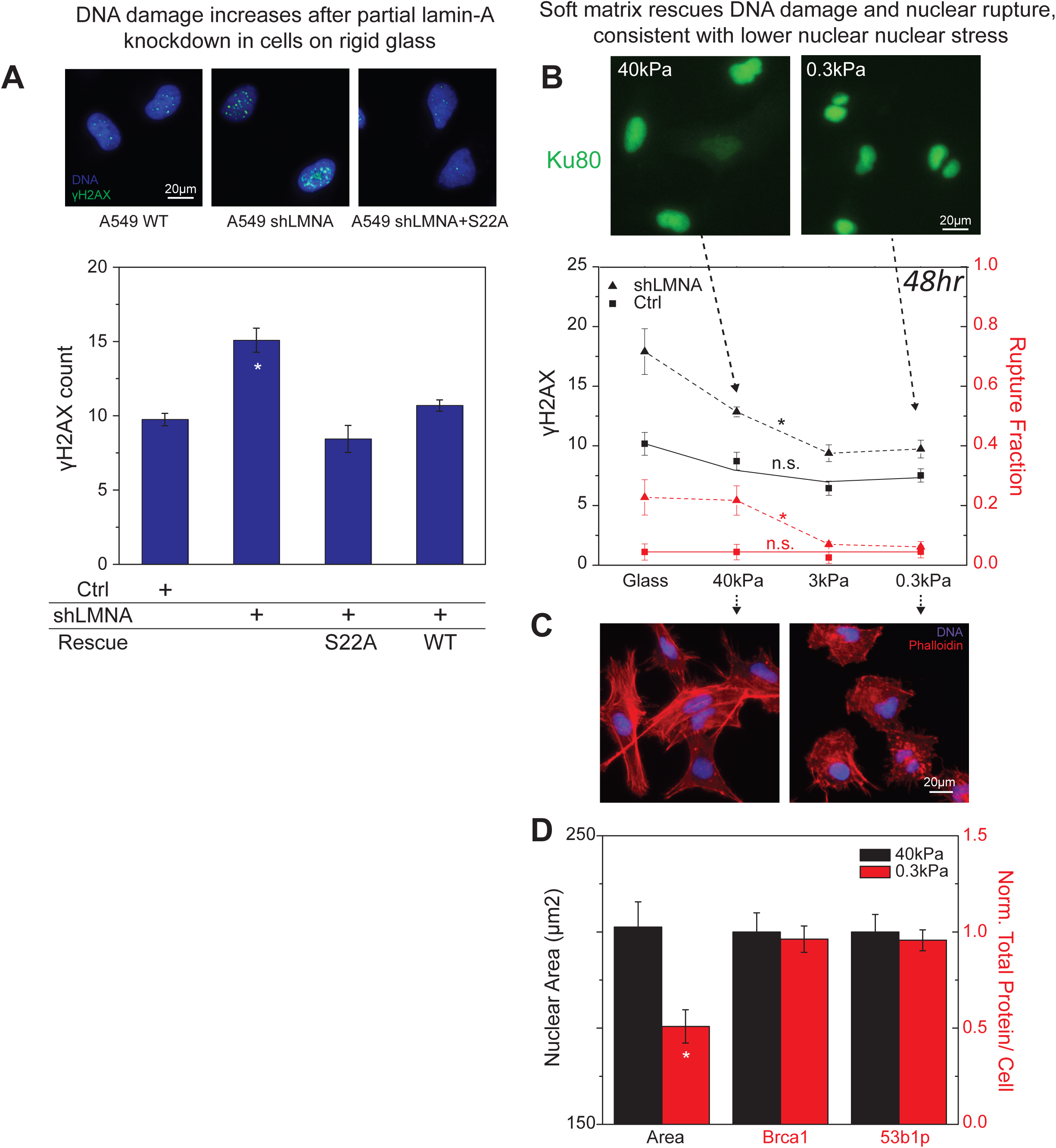
Growth of cells on soft matrix relaxes cytoskeletal stress on the nucleus, suppresses the loss of DNA repair factors, and thereby limits DNA damage. (A) A549 Ctrl, S22A and WT rescued cells show lower γH2AX foci number than shLMNA cells on rigid plastic. (>100 cells per condition, average ± SEM. *P<0.05 to Ctrl) (B) Soft matrix rescues nuclear rupture and DNA damage after 48 hrs culture. (>100 cells per condition, n=2 experiment, average ± SEM. *P<0.05) (C) shLMNA cells on 40kPa gel have more stress fibers (Phalloidin) than cells on 0.3 kPa gel. (D) Nuclear area of shLMNA cells shrinks slightly on soft matrix, but total 53BP1 and BRCA1 protein levels per cell stay constant. (>100 cells per condition, n=2 experiment, average ± SEM. *P<0.05 to 40kPa)

Soft gels are a better mimic of soft tissues^18^ and reportedly decrease nuclear rupture frequency of a Lamin-A progeroid fibroblast line compared to growth on stiff gels^19^. However, previous studies with gels do not seem to have examined a simple knockdown of Lamin-A nor any effect on repair factors. Mis-localization of Ku80 into the cytoplasm of Lamin-A knockdown cells is indeed reduced 5-10 fold in cells grown on soft gel substrates compared to knockdown cells on both stiff gels and conventional rigid glass coverslips (Fig 4B). Crucially, γH2AX foci counts also decrease 5-10 fold for knockdown cells on soft gels (Fig 4B). Control cells with normal levels of Lamin-A show lower overall levels of mis-localization and DNA damage in cells on stiff gels, and soft gels lead to weak, non-significant decreases. Repair factor mis-localization and enhanced DNA damage for knockdown cells is thus partially recovered by the soft gels.

Cells on stiff gels spread well and maintain an abundance of long stress fibers compared to cells on soft gels (Fig 4C). Such spreading stresses the nucleus and tends to increase its projected area in both A549 cells and human MSCs^18^; the A549 cells with knockdown of Lamin-A likewise show a reduction in nuclear area on soft gels (Fig 4D). Importantly, BRCA1 and 53BP1 protein levels in Lamin-A knockdown cells remain constant over the 48 hours of culture on soft and stiff matrix. Increased DNA damage in cells on rigid substrates relative to soft substrates (Fig 4B) is therefore consistent with repair factor mis-localization after rupture of a highly stressed nucleus.

### Lamin-A depletion increases DNA damage before and after constricted migration

Constricted migration causes nuclear rupture with mis-localization of repair factors (53BP1 in Fig.1B,C; and Ku80 and BRCA1 in recent studies^9, 10^) and increased DNA damage (Fig.1 B; *ibid*), while 2D cultures on rigid substrates of Lamin-A knockdown cells cause similar effects (Fig 4B) with eventual (>> 48 hrs), permanent loss of at least some repair factors – particularly BRCA1 (Fig 2C, 3C). Lamin-A knockdown cells indeed show more nuclear blebs than Ctrl cells before migration through a rigid transwell filter with 3 μm pores (Fig 5A) and also exhibit more DNA damage (Fig 5B). However, nearly all cells of either type show long-lived blebs after the 24 hrs of migration (Fig 5A), which suggests that migration induced rupture is not a distinctive aspect of mechanism. DNA damage is nonetheless much higher for Lamin-A knockdown cells after migration, and γH2AX once again exhibits a nucleoplasmic distribution rather than any evident enrichment at a bleb or near the NE (Fig 5B). Low levels of BRCA1 (and BRCA2) are known to slow the exponential decay by many hours in γH2AX-marked damage that spikes after a brief pulse of ionizing radiation^20^. Constricted migration is thus a similar perturbation that occurs over a few hours (Fig 5C) and likewise causes DNA damage that decays over time as repair factors recover^10^.

**Fig. 5.**
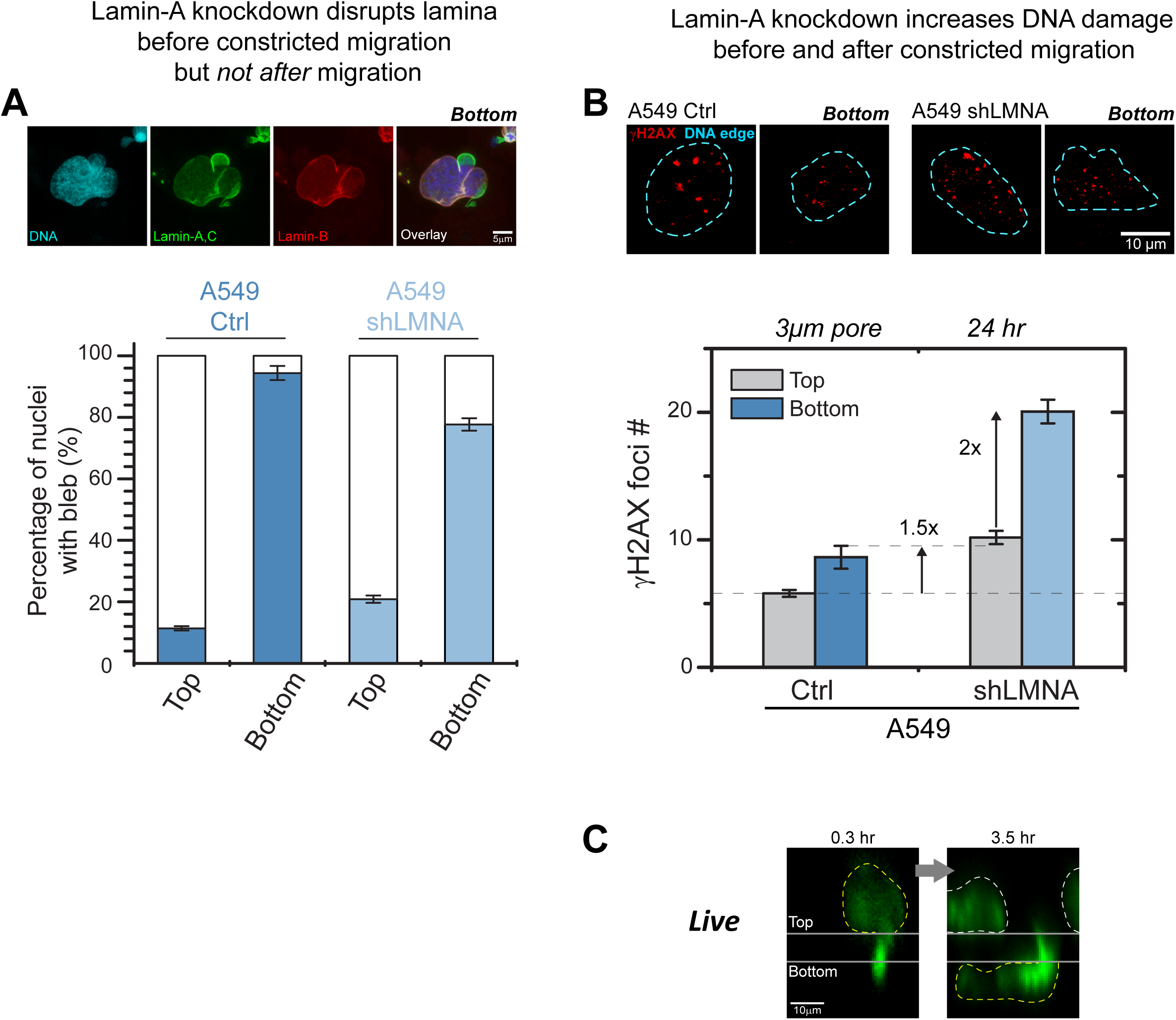
DNA DSBs is largely enhanced in shLMNA cells comparing to Ctrl cells post-migration, but bleb fraction is not enhanced. (A) Lamin-A knockdown disrupts lamina before constricted migration but not after constricted migration. Elevated blebs formation in both Ctrl and shLMNA A549 cells was observed after migration through 3 μm pores, but shLMNA cells do not show more bleb fraction than Ctrl. (B) γH2AX foci are counted using ImageJ algorithm and foci distribution is shown. (n = 51-578 per condition per experiment, n=4-5 experiments, average ± SEM.) (C) Live imaging indicates A549 cells spend 3 hours migrating through 3μm pores.

Despite many similarities between nuclear rupture of lamin-A compromised cells grown on rigid substrates and wildtype cells in constricted migration, there are clearly some differences that extend beyond stable changes in repair factors such as 53BP1 and BRCA1 (Fig 3). For example, constricted migration causes most cells to exhibit long lived nuclear blebs that are more transient and less frequent for lamin-A defective cells (Fig 2F; and past work^7,5^). Cells that migrate to the sparse but rigid bottom of a transwell will spread and flatten the nucleus (Fig 5C), thereby stressing the nuclear lamina. Perhaps the lamina rupture that occurs only with small pores^8,10^ is then stressed sufficiently to delay lamina remodeling and repair. Such issues highlight the need to more deeply understand the effects of microenvironment on nuclear stress and mechanosensing^18^.

## Materials and methods

### Cell culture

U2OS – human osteosarcoma cell line and A549 - human lung carcinoma cell line, were cultured in DMEM high glucose media and F12 media (Gibco, Life Technologies) respectively, supplemented with 10% FBS and 1% penicillin/streptomycin (Sigma-Aldrich) under 37°C and 5% CO_2_ as recommended by ATCC.

### Transwell migration

Cells were seeded at 300,000 cells/cm^2^ onto the top side of the Transwell membrane (Corning) and allowed them to migrate in normal culture condition with serum for 24 hours. To isolate the cells from the membrane, cells were detached from the membrane by using 0.05% Trypsin-EDTA (Gibco, Life Technologies).

If the isolated cells were to undergo another transwell migration, they were expanded for a week to reach the required number of cells.

### Immunostaining and imaging

Cells were fixed in 4% formaldehyde (Sigma) for 15 minutes, followed by permeabilization by 0.25% Triton-X (Sigma) for 10 minutes, blocked by 5% BSA (Sigma) and overnight incubation in various primary antibodies: lamin-A/C (Santa Cruz and Cell Signaling), Lamin-B (Santa Cruz), γH2AX (Millipore), 53BP1 (Novus), Ku80 (Cell Signaling), BRCA1 (Santa Cruz), RPA1 (Santa Cruz), phosphorylated S1981 ATM (Abcam). The cells were then incubated in secondary antibodies (ThermoFisher) for 1.5 hours, and their nuclei were stained with 8μM Hoechst 33342 (ThermoFisher) for 15 minutes.

For Transwell staining, the entire Transwell membrane was fix in 4% formaldehyde, followed by normal IF procedures described above. When mounting is involved, Prolong Gold antifade reagent was used (Invitrogen, Life Technologies). Epifluorescence imaging was performed using Olympus IX71, with a digital EMCCD camera (Cascade 512B, Photometrics) and a 40×/0.6 NA objective. Confocal imaging was done in Leica TCS SP8 system, by either 63×/1.4 NA oil-immersion or 40×/1.2 NA water-immersion objectives. Various image quantification and processing were done with ImageJ.

### Synthesis of soft and stiff polyacrylamide gels

Round glass coverslips (18mm, Fisher Scientific) were cleaned in boiling ethanol and RCA solution (H2O:H2O2:NH4OH=2:1:1 in volume) for 10min each, and then functionalized in ATCS solution (Chloroform with 0.1% allytrichlorosilane(Sigma) and 0.1% triethylamine(Sigma)) for an hour. Fresh precursor solution for 0.3kPa soft gels (3% acrylamide + 0.07% bis-arylamide in DI water) and 40kPa stiff gels (10% acrylamide + 0.3% bis-acrylamide in DI water) were prepared respectively. Afterwards, 0.1% N,N,N,N-tetramethylethylenediamine (Sigma) and 1% ammonium persulphate(Sigma) were added to each precursor solution and pipetted 20ul of this mixture onto each coverslip to allow gel polymerization. To coat collagen-I on the gel, crosslinker sulpho-sanpah (50ug/ml in 50mM HEPES, G-Biosciences) was applied to cover the whole gel surface and photoactivated under 365nm UV light for 7min. Excess amount of sulpho-sanpah were washed after UV and collagen-I solution (100¦Ìg/ml in 50mM HEPES) was added for coating. The coating process usually takes overnight on shaker at room temperature to ensure saturation.

### Protein modulation in U2OS cells

All siRNAs used in this study were purchased from Dharmacon. U2OS cells were passaged 24 hours prior to transfection. A complex of siRNA pool [25 nM siLMNA, siCtrl] and 1 μg/mL Lipofectamine 2000 was prepared according to the manufacturer’s instructions and incubated for 24 hours in high corresponding media with 10% FBS. Knockdown efficiency was determined by Western blot following standard methods.

### Viral transduction

In order to establish an A549 shLaminA line, lenti-vector was added together with 8 μg/mL polybrene (Milipore) and 200 μL of full culture media for 2 days. The knockdown level was confirmed by Western blot following standard methods

### Statistical analysis

A ‘two-tailed’ Student’s t-test was used to calculate the statistical significance of the observed differences. Microsoft Excel 2013 was used for the calculations. In all cases, differences were considered statistically significant when P<0.05. Unless otherwise indicated, values represent mean±s.e.m.

